# Limits of chromosome compaction by loop-extruding motors

**DOI:** 10.1101/476424

**Authors:** Edward J. Banigan, Leonid A. Mirny

## Abstract

During mitosis, human chromosomes are linearly compacted ~ 1000-fold by loop-extruding motors. Recent experiments have shown that condensins extrude DNA loops, but in a “one-sided” manner. We explore whether one-sided extrusion can compact chromosomes by developing a mean-field model for polymer compaction by motors that actively extrude loops and turnover. The model establishes an upper bound of only ~ 10-fold for compaction by one-sided extrusion. Thus, it cannot be the sole mechanism of chromosome compaction. However, other, effectively two-sided mechanisms can achieve sufficient compaction.

---

Chromosomes are exceedingly long polymers of chromatin, *i.e.*, DNA and associated proteins. Each chromosome in a human cell is comprised of ~ 1 mm of chromatin, which is dynamically reorganized throughout the cell cycle. During mitosis, chromosomes are linearly compacted ~ 1000-fold forming ~ 1 − 5 *µ*m-long cylindrical shapes. This compaction is achieved by the formation of an array of loops [1–8]. While proteins essential for this process are known, the physical mechanisms underlying this dramatic compaction are not fully understood.

The main molecular driver of compaction is condensin, a ring-like protein complex [9–11]. It has been hypothesized that condensin and other SMC (Structural Maintenance of Chromosomes) complexes are molecular motors that actively extrude chromatin loops [12–14]. These loop-extruding factors (LEFs) load onto the chromatin fiber and progressively form larger loops by reeling chromatin into the loops (Fig. 1(a)). Models suggest that loop extrusion can dynamically generate various chromosome structures [15–19], including linearly compacted mitotic chromosomes. Experiments support loop extrusion as a common phenomenon in living cells [7, 8, 20–30], but this motor activity had not been directly observed until recently.

**FIG. 1.**
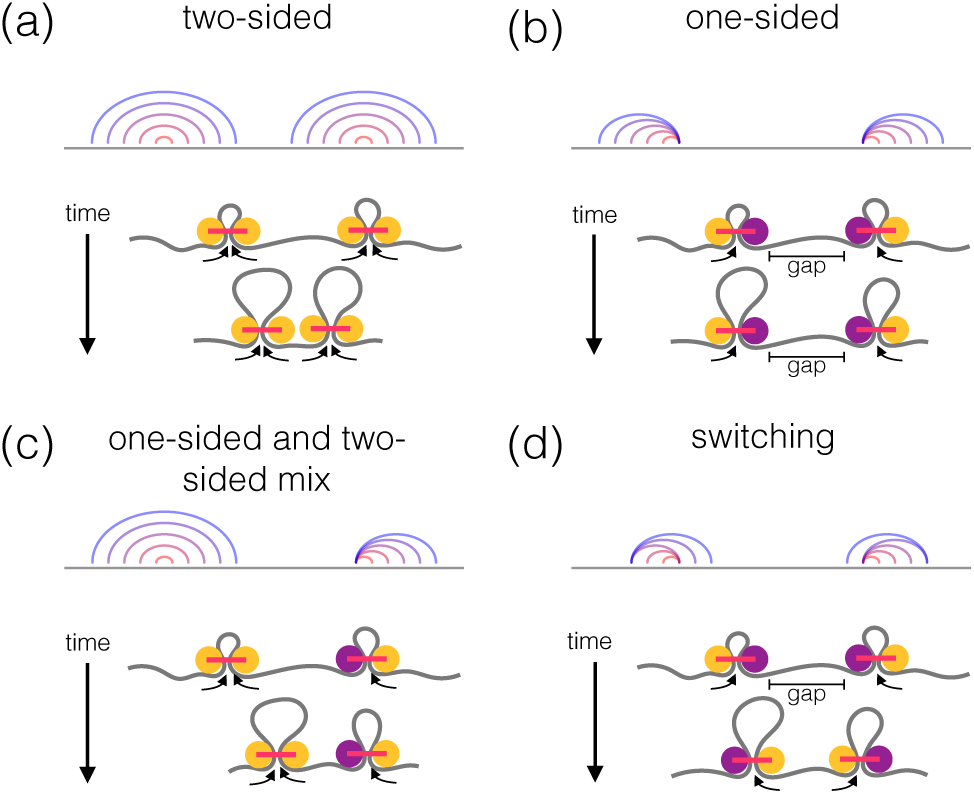
(a) *Top*: Arch diagram showing positions of two “two-sided” loop-extruding factors (LEFs) at different times, with time indicated by color from red (early) to blue (late). *Bottom*: Drawing of two-sided LEFs (yellow and pink) translocating along chromatin/DNA (gray) and progressively growing loops. (b) Arch diagram with time progression of “one-sided” LEF positions and drawing of one-sided LEFs, each with its inactive head shown in purple. LEFs in the depicted configuration leave an unextruded gap. (c) A mixture of one- and two-sided LEFs. (d) One-sided LEFs that switch which side extrudes can eliminate initially unextruded gaps if at least one of the LEFs switches.

Recent *in vitro* single-molecule experiments demonstrated that yeast condensins actively extrude DNA loops in an ATP-dependent manner [31], supporting the loop extrusion mechanism. However, loop extrusion in these experiments was “one-sided” so that DNA on only one side of the condensin was reeled into the loop (Fig. 1(b)). This conflicts with previous models, which assumed that each LEF consists of two connected motors that reel chromatin from both sides, and thus performs “two-sided” extrusion (Fig. 1(a)). This raises the question of whether one-sided extrusion can also generate the chromosome structures attributed to loop extrusion, particularly linearly compacted mammalian chromosomes.

To investigate the ability of condensin to compact chromosomes, we develop a mean-field theoretical model for active LEFs that transiently load onto DNA and extrude loops in a directed manner. We consider the cases of two-sided LEFs (Fig. 1(a)), one-sided LEFs (Fig. 1(b)), a mix of one- and two-sided LEFs (Fig. 1(c)), and one-sided LEFs that can switch which side extrudes (Fig. 1(d)). We predict upper limits for compaction in these scenarios, which we confirm with stochastic simulations. Surprisingly, one-sided loop extrusion can achieve only up to 10-fold compaction, even with LEF turnover. While this may be sufficient for yeast chromosomes [32], it cannot be the mechanism underlying ~ 1000-fold mammalian chromosome compaction.

We model LEFs as two-headed complexes with either one or two active heads (Fig. 1(a),(b)). Each active head directionally translocates along chromatin/DNA at speed *v* in one dimension, as observed *in vitro* [31, 33]. For one-sided extrusion, one head is inactive and stationary [31], so only the active head reels chromatin into the extruded loop, which grows at speed *v*. For two-sided extrusion, both heads are active, and the loop grows at speed 2*v*. To model condensin loading and unloading dynamics [33– 36], LEFs stochastically bind chromatin at rate *k_b_* and unbind at rate *k_u_*, leading to a steady-state average of *N_b_* = *k_b_N/*(*k_u_* + *k_b_*) of polymer-bound LEFs, where *N* is the total number of LEFs in the system. A LEF may load within an existing loop, thus forming a *child* loop that either splits or reinforces the *parent* loop (Fig. 2(a)). As shown for two-sided extrusion [16], two length scales control the dynamics of the system: LEF processivity, *λ* = 2*v/k_u_*, *i.e.*, the average size of a loop formed by an unobstructed LEF, and the average separation, *d* = *L/N_b_*, of LEFs on the polymer. For *d ≫ λ*, LEFs are sparse so that they do not obstruct each other and rarely nest, and the polymer is not significantly compacted. In contrast, *d ≪ λ* is a dense regime in which most of the polymer is extruded into reinforced loops, which yields high (*>* 500-fold) linear compaction. For two-sided extrusion in this regime, the entire polymer can be extruded into loops because extrusion eliminates all gaps between loops. One-sided extrusion, however, unavoidably leaves gaps between LEFs leading to incomplete compaction.

**FIG. 2.**
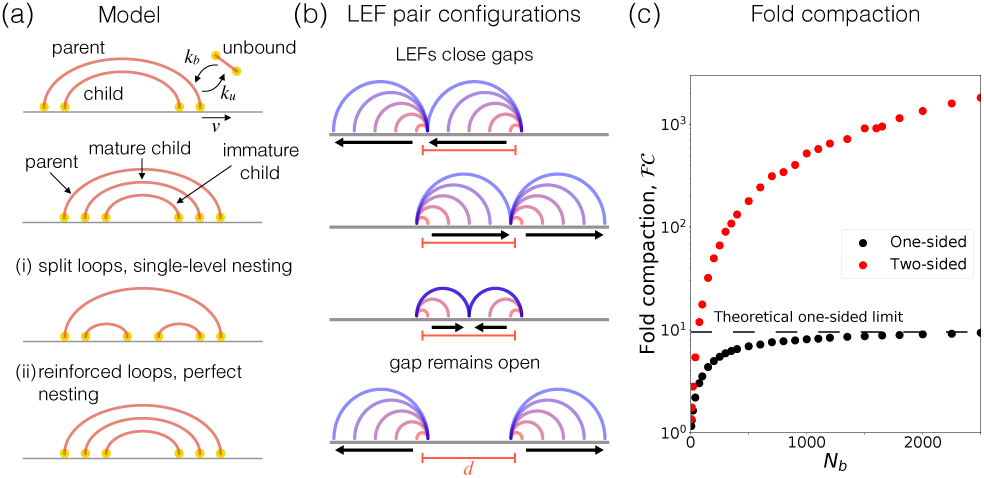
(a) *Top half*: A parent LEF is associated with chromatin at the base of an extruded loop and a child LEF is associated within the loop. A bound LEF can unbind chromatin to become an unbound LEF, and an unbound LEF can bind to become a parent or child. A “mature” child LEF will become a parent LEF if the current parent LEF unbinds, while a more deeply nested “immature” child LEF would remain a child. *Bottom half*: Loop nesting architectures. With (*i*) single-level nesting, child LEFs split parent loops, while with (*ii*) perfect, nesting, child LEFs reinforce the parent loops. (b) Possible configurations of pairs of adjacent one-sided LEFs, each shown at different times indicated by color from red (early) to blue (late). Side of extrusion (and direction of motor translocation) is indicated by arrows. One configuration – where active LEF subunits translocate away from each other by extruding chromatin from opposite sides of the polymer – leaves a gap. (c) Simulation results for fold compaction,, by one-sided extrusion (black) and two-sided extrusion (red), shown for varying number of bound LEFs, *N_b_*.

To examine compaction by one-sided extrusion, we consider possible orientations of neighboring one-sided LEFs. In three of four possible configurations, one-sided LEFs can close the gap between them if they remain bound to the polymer for a sufficiently long time. In the remaining configuration, adjacent LEFs necessarily leave a gap between them because the intervening polymer is not extruded by either LEF (Fig. 2(b)). Thus, in the dense regime (*λ/d* ≫ 1), where extrusion is much faster than exchange kinetics, one-sided LEFs segment chromatin into *N_p_* parent loops and *N_p_/*4 gaps between the loops.

To calculate the maximum fold compaction 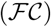 achievable by one-sided LEFs, we consider the maximum compacted fraction, *f*, defined as the fraction of the polymer extruded into loops. These quantities are related as 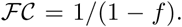. Hence, even a modest fraction, 1 − *f*, of unextruded polymer in gaps strictly limits the degree of linear compaction of the polymer.

We start by calculating mean loop and gap sizes. The LEFs segment chromatin of length *L* into parent loops of mean size:

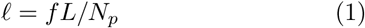

and gaps with mean size equal to the mean distance between the loading sites of two adjacent LEFs (Fig. 2(b)):

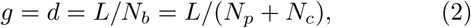

where *N_c_* is the number of child LEFs. A key assumption of the model is the existence of these unextruded gaps. As explained above, we expect gaps to occur for one of four LEF configurations (Fig. 2(b)), so a total of *N_p_* parent loops and *N_p_/*4 gaps cover the whole polymer:

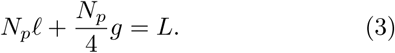

Combining Eqs.      (1)-(3), we obtain the compacted fraction that has been extruded into loops:

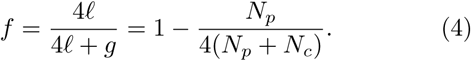

This indicates that significant lengthwise compaction by one-sided LEFs requires a large number of child LEFs per parent (*e.g.*, *f* = 0.99, or 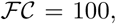, requires *N_c_* ≈ 25*N_p_*).

To explore whether this picture can be altered by LEF turnover, we determine the steady-state kinetics of unbound, parent, and child LEFs. A new parent is formed if one of the *N_u_* unbound LEFs binds to a chromosomal position that is not within an extruded loop. Alternatively, an unbound LEF becomes a child LEF if it binds to a site within an extruded loop (Fig. 2(a)). Thus, upon binding, an unbound LEF becomes a child LEF with probability *f* or a parent LEF with probability 1 − *f*. Upon unbinding of a parent LEF, child LEFs previously nested within the parent loop may become parents if they are “mature,” *i.e.*, not nested within any loop other than the parent (Fig. 2(a)). The average number of mature children per parent, denoted by *α*, depends on the loop nesting architecture. LEF kinetics are therefore described by:

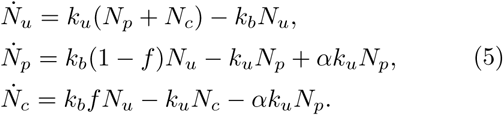

Solving these equations in steady state yields a relation between the numbers of child and parent loops:

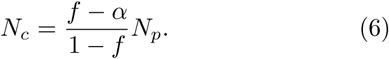

This equation fixes the ratio of child LEFs to parent LEFs for each possible *f*. Moreover, using Eq. (6) to substitute for *N_c_* in Eq. (4), the mean number of mature child loops per parent loop in steady state is *α* = 3*/*4. However, to completely solve the system and determine the maximum compaction *f* from Eq. (4), we still must determine the relationship between *N_p_* and *N_c_*.

To find the maximum possible compaction we consider two limiting loop architectures: *i*) single-level nesting in which all child LEFs are mature, so that child LEFs only split parent loops, and *ii*) perfect nesting in which each parent loop is reinforced by at most one mature child LEF, and each child LEF is reinforced by at most one child (Fig. 2(a)).

With single-level nesting, *α* = *N_c_/N_p_* = 3*/*4, and each child LEF becomes a parent when its parent LEF unbinds (Fig. 2(a)i). From Eq. (4) the maximum compaction is:

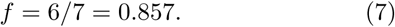

We also find LEFs per parent loop, (*N_p_* + *N_c_*)*/N_p_* = 7/4 and loop sizes, *f* = *f L/N_p_* = 3*d/*2.
With perfect LEF nesting, each parent loop has at most one mature child (Fig. 2(a)ii). We calculate the probability that a parent loop does not have a child in the limit of large *N_p_* and *N_c_*:

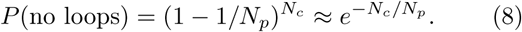

The probability that a parent loop contains a child loop is thus *α* = 1 *− e^−Nc /Np^* = 3*/*4, and we find:

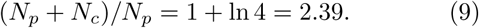

Combining Eqs. (4) and (9), we find the compaction limit:

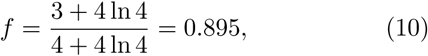

or 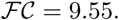. Additionally, the mean loop size is:

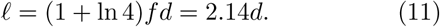

In the dense regime (*λ/d ≫* 1), the mean distance between LEFs, *d* = *L/N*, controls the scaling behavior of the system. For instance, increasing the number of LEFs, *N*, decreases the gap size as 1*/N*, but also increases then number of gaps as *N*, fixing the maximum compacted fraction *f* [37]. Thus, both cases achieve at most ~ 10-fold compaction, along with small loops and little LEF nesting.

Importantly, Eq. (10) gives the maximum degree of chromatin compaction achievable by one-sided loop extrusion. To see that perfect nesting gives the upper bound to the maximum compaction (Eq. (10)) while single-level nesting corresponds to the lower bound (Eq. (7)), we rewrite Eq. (6) as:

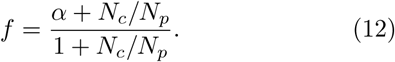

This is an increasing function of *N_c_/N_p_* for *N_c_/N_p_ >* 0. The two cases are the lower and upper bounds for *N_c_/N_p_* and thus, the two extremes of possible loop architectures. They correspond to the bounds for the compaction limit; the compaction limits of all other architectures fall within the range defined by these extremes. Thus, we predict that chromosomes cannot be linearly compacted by more than an average of 10-fold for any loop architecture generated by one-sided LEFs with independent kinetics.

To test the predictions of the mean-field theory, we use stochastic simulations to measure polymer compaction by one-sided LEFs. Adapting the simulation model in [16], we simulate *N* one-sided LEFs that extrude loops at speed *v* and have stochastic kinetics described by Eq. (5). The simulations confirm that the mean-field theory with perfect nesting accurately predicts the fraction compacted, *f*; for large *λ/d*, *f* = 0.895 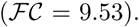. in simulations (see large *N_b_* in Fig. 2(c), *ϕ* = 0 in Fig. 3(a), and Eq. (10)). Moreover, the simulations illustrate the drastic decrease in fold-compaction achievable with one-sided LEFs as compared with two-sided LEFs (Fig. 2(c)). The simulations also confirm other predictions of the theoretical model. Given the possible configurations of LEFs (Fig. 2(b)), we expect one gap of size *g* = *d*, per four loops. In accord with the theory, we observe *N_g_/N_p_* = 0.22, and mean gap size, *g ≈ d* in the simulations (Fig. S1 [38]). Moreover, consistently with the theory (*α* = 3*/*4), the number of mature child LEFs per parent LEF in steady-state simulations is *α* = 0.76. We also observe (*N_p_* +*N_c_*)*/N_p_* = 2.13 LEFs per loop (Eq. (9)) and a mean loop size of *f* = 1.90*d* (Eq. (11)). Differences in (*N_p_* + *N_c_*)*/N_p_* and *f* between simulation and theory are due to the theoretical assumption of perfectly nested loops, which is violated by 18% of parent loops. Deviations in (*N_p_* + *N_c_*)*/N_p_* and *l* offset each other when computing *f*; the theory predicts fewer but larger loops than observed in simulations. Altogether, the simulation results indicate that the mean-field theory accurately describes LEF dynamics, loop architecture, and maximum polymer compaction in the dense regime (*λ/d ≫* 1).

**FIG. 3.**
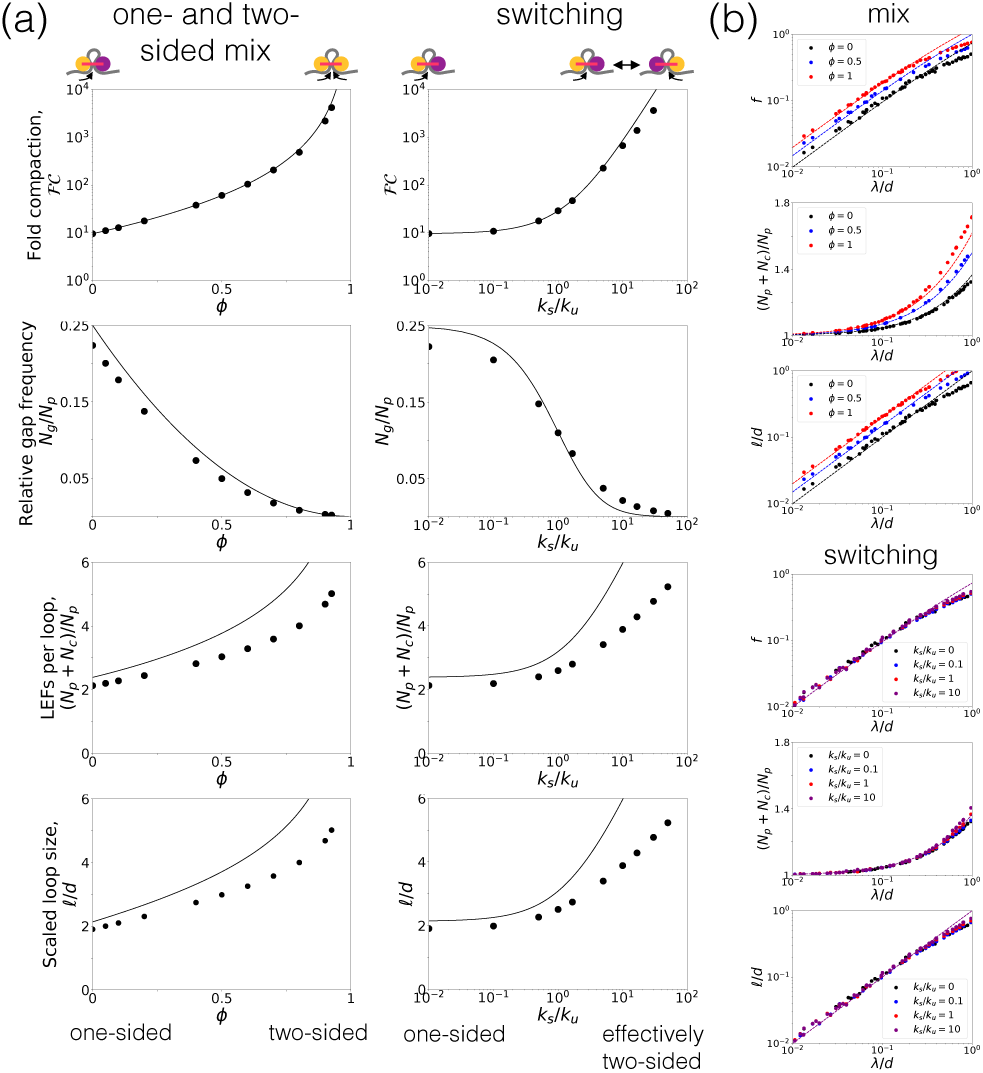
(a) Simulation (points) and theory (lines) in the dense *λ/d* ≫ 1 regime for models with a mix of one- and two-sided LEFs (left) and switching (right). From top to bottom: Fold compaction 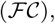, relative gap frequency (*N_g_/N_p_*), LEFs per loop ((*N_p_*+ *N_c_*)*/N_p_*), and scaled loop sizes (*R/d*). Note *ϕ* = 0 (and *k_s_/k_u_*= 0) for pure one-sided extrusion and *ϕ* = 1 and (*k_s_/k_u_* →) for (effectively) two-sided extrusion. (b) Fraction compacted (*f*), LEFs per loop, and scaled loop sizes in the sparse *λ/d <* 1 regime.

We have found that one-sided LEFs with the minimal properties observed in single-molecule experiments [31, 33] are unable to achieve the dramatic polymer compaction needed to package mammalian mitotic chromosomes *in vivo* [4, 5]. To explore how one-sided loop extrusion may nonetheless facilitate mitotic chromosome compaction, we extend the theory to determine which loop extrusion variants can fully compact mitotic chromosomes.

We consider a model in which a fraction *ϕ* of the LEFs are two-sided and 1 − *ϕ* are one-sided (Fig. 1(c)). Since two-sided LEFs can close all gaps, there is one configuration of LEFs that leaves a gap out of nine possible configurations. The frequency with which this configuration appears is *N_g_/N_p_* = (1 *− ϕ*)^2^/4. The equation relating loops and gaps (previously, Eq. (3)) becomes:

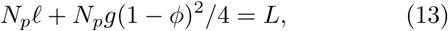

so that the compaction fraction in this model is given by:

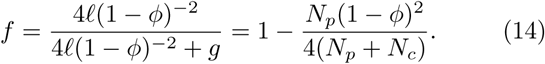

High compaction can be achieved with little LEF nesting when enough two-sided LEFs are present. For *λ/d* ≫ 1, LEFs form reinforced loops [16], so we assume perfect nesting and note that α = (3 + 2ϕ − ϕ^2^/4, to find:

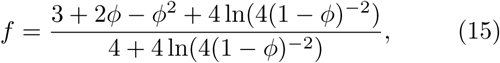

which is plotted in Fig. 3(a). The theory also predicts the LEFs per loop (*N*_*p*_ + *N*_*c*_) = *N*_*p*_, mean loop size *l*, and number of gaps per loop *N*_*g*_/*N*_*p*_ observed in simulations (Fig. 3(a), left). The simulations show that the mean-field theory based on as a predicted number of gaps per loop accurately captures the maximum compaction achievable by mixtures of one- and two-sided LEFs (Fig. 3(a)).

According to Eq. (15), a 1:1 mix of one- and two-sided LEFs yields *f* = 0.983, or 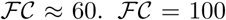 requires a fraction of ϕ = 0.59 two-sided LEFs and 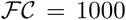 requires ϕ = 0.84. Thus, even a modest fraction of one-sided LEFs can perturb mitotic chromosome compaction. As expected for *λ/d* ≫ 1, pure two-sided extrusion (ϕ = 1) yields 100% compaction. Thus, to eliminate unextruded gaps and fully compact chromosomes, a large majority of the LEFs must be two-sided, which contrasts with observations from *in vitro* experiments.

Therefore, we consider an alternative model in which one-sided LEFs close gaps without simultaneously extruding from both sides. Here, LEFs are instantaneously one-sided, but each LEF stochastically switches which side actively extrudes at rate *k_s_* (Fig. 1(d)). In principle, switching could occur due to exchange of subunits of the SMC complex while the complex remains loaded [35], alterations to solution conditions [31], or post-translational or genetic modifications [39, 40].

Compaction is limited by the degree to which the LEFs act in a purely one-sided manner. LEFs that switch before unloading close adjacent gaps, regardless of the orientation of the adjacent LEFs, whereas LEFs that do not switch are effectively one-sided. We calculate the fraction, *n*_2_, of one-sided LEFs that switch before unloading by solving

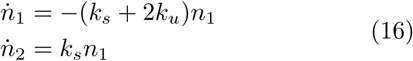

and finding *n*_2_ ≡ lim_*t→∞*_ n_2_. In the 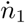 equation, 2*k_u_* is used because in order for a LEF in a LEF pair to be effectively two-sided, it must switch before either it or its neighbor unbinds. The fraction of effectively two-sided LEFs is:

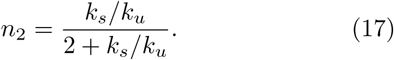

This fraction only depends on *k_s_/k_u_*, the rate of switching compared to the unloading rate. For fast switching, *n*_2 → 1_.

By substituting *n*_2_ for the fraction, *ϕ*, of two-sided LEFs into Eq. (15), we find the compaction limit as a function of *k_s_/k_u_*:

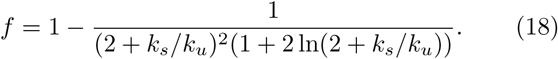

Again, the theory predicts values for the compaction limit, gaps per loop, LEFs per loop, and mean loop size that agree with simulation observations (Fig. 3(a), right). Eq. (18) indicates that a fast, but physiologically plausible, switching rate is required to achieve robust compaction. For example, 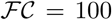 requires *k_s_/k_u_* = 2.9, while 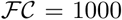 requires *k_s_/k_u_* = 10.8. The unloading rate, *k_u_*, is observed to be of order 0.1 min^*−*1^ [33–36, 41– 43], so the expected switching rate is of order 1 min^*−*1^. However, switching was not observed over 1-5 minutes of observations in recent *in vitro* single-molecule experiments [31]. Thus, LEF switching would require some as yet unknown *in vivo* factor or condition. Nonetheless, the theory and simulations indicate that effectively two-sided extrusion is required for 1000-fold mitotic chromosome compaction.

The theory can be extended to another physiologically important regime that is relevant to the SMC complex cohesin. While have explored the dense (*λ/d* ≫ 1) regime relevant to condensin-driven compaction of mitotic chromosomes, organization of interphase topologically associated domains (TADs) is likely driven by loop-extruding cohesins with *λ/d* ≲ 1 [18, 19, 25, 26]. Thus, we explore the sparse *λ/d <* 1 regime, in which LEFs leave unextruded gaps on both sides because LEF processivity is smaller than the distance between LEFs.

To compute compaction, *f*, note that each loop grows unimpeded and loop size is the processivity, so *l ≈ λ*(1 − *ϕ*) + 2*λϕ* (with *λ* = *v/k_u_* and *ϕ* = 0 for the switching model since only one subunit is active at any time). We substitute for *α* in Eq. (6) and use *f* = *f L/N_p_* = *f d*(*N_p_* + *N_c_*)*/N_p_*. For both perfect nesting and single-level nesting, with small *λ/d* and *f*, we find [38]:

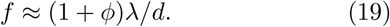

Simulations confirm this theoretical result (Fig. 3(b)). Thus, for *λ* ≪ 1, compaction is linear in *λ/d* and loop nesting is unimportant because it is rare. This is true for both one- and two-sided extrusion in the sparse regime, but whether one-sided extrusion can organize chromosome structures such as interphase TADs in eukaryotes [20–26] remains unresolved.

Together, the theory and simulations show that the linear compaction of mitotic chromosomes is dependent on the ability of LEFs to eliminate uncompacted gaps between extruded loops. Thus, one-sided loop extrusion alone is unable to robustly compact chromosomes by more than 10-fold (Figs. 2(c) and 3(a)) because it in-evitably leaves one uncompacted gap for every four loops (Fig. 2(b)). These gaps remain in the steady state of dynamically exchanging LEFs. Moreover, these gaps cannot be reliably eliminated if the “safety belt” that anchors each one-sided LEF [31, 39] instead freely diffuses along DNA because loop growth would be inhibited by configurational entropy [44]. 10-fold linear compaction may be sufficient for yeast chromosomes [32], but it is inconsistent with *in vivo* observations of *>* 100-fold linear compaction of mammalian chromosomes [4, 5].

A possible explanation is that yeast condensins are one-sided extruders as observed *in vitro* [31], while mammalian condensins are two-sided. Yeast chromosomes are of order 1 megabase pair (Mbp), so they may require less compaction than human chromosomes, which are of order 100 Mbp. To see this, note 1 bp = 0.34 nm and DNA has persistence length *f_p_* = 50 nm (note that a similar argument holds if we consider a chromatin fiber with nucleosomes [45]). We expect uncompacted 3D chromo-some lengths of:

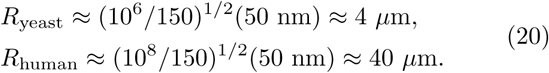

Thus, 10-fold linear com_*√*_<u>pa</u>ction which may reduce the 3D size by a factor of 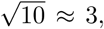, produces micron-sized yeast chromosomes, but 10-micron human chromosomes. Human chromosomes need *>* 100-fold linear compaction for 3D size of order microns.

A complementary hypothesis is that condensin performs two-sided extrusion facilitated by *in vivo* factors or conditions. A plausible mechanism of effectively two-sided extrusion is directional switching, which could be induced by subunit exchange [35], ambient conditions (*e.g.*, concentration or ionic composition) [31], or protein modifications [39, 40]. Another possibility is that LEFs extrude asymmetrically with speeds *v*_1_ *> v*_2_, in which case we would observe effectively two-sided extrusion if the slow side is fast enough to close gaps before disassociation (*i.e.*, *v*_2_*/k_u_ ≫ d*) [44]. Alternatively, LEFs could be effectively two-sided if condensins dimerize or otherwise cooperate to form two-sided complexes. Oligomerization is consistent with experimental observations for both condensin [46–48] and cohesin [49–51].

Our theoretical model couples LEF loading and unloading kinetics with mean-field assumptions for the resulting loops. In this model, we find that linear compaction by one-sided loop extrusion is strictly limited. The model indicates that robust linear compaction for any type of LEF relies on minimizing the frequency of gaps of uncompacted chromatin (Eqs. (3) and (13)). Thus, biological mechanisms that favor cooperativity or effectively two-sided loop extrusion are likely necessary to achieve robust compaction of mitotic chromosomes. In particular, the model predicts that mammalian condensin complexes perform effectively two-sided loop extrusion under *in vivo* conditions.

## ACKNOWLEDGMENTS

We thank John Marko, Anton Goloborodko, Cees Dekker, Kikuë Tachibana, and Maxim Imakaev for helpful discussions and Johannes Nuebler for critically reading the manuscript. This work was supported by the NSF Physics of Living Systems (15049420) and by the Center for 3D Structure and Physics of the Genome of NIH 4DN Consortium (DK107980).

