## Supplementary material for "Limits of chromosome compaction by loop-extruding motors"

### Limits of chromosome compaction by loop-extruding motors: Supplementary information

(Dated: November 21, 2018)

#### COMPACTION IN THE SPARSE REGIME

The theory can be extended to the sparse $\lambda/d < 1$ regime, in which LEFs leave unextruded gaps on both sides because LEF processivity is smaller than the distance between LEFs. To compute the compaction, $f$, in this regime, note that each loop grows unimpeded and loop size is the processivity, so

$$\ell \approx \lambda(1-\phi) + 2\lambda\phi. \tag{1}$$

Here, note that $\phi = 0$ for the switching model since only one head is active at any time. To find compaction, we substitute for $\alpha$ in Eq. (6) in the main text and use

$$\ell = fL/N_p = fd(N_p + N_c)/N_p. \tag{2}$$

With single-level loop nesting, $\alpha = N_c/N_p$, so $\ell = \lambda(1+\phi) = fd(1-f/2)^{-1}$, which yields:

$$f = \frac{2(1+\phi)\lambda/d}{2+(1+\phi)\lambda/d} \approx (1+\phi)\lambda/d. \tag{3}$$

With perfect nesting, $\alpha = 1 - e^{-N_c/N_p}$, so $\lambda(1+\phi) = fdW(e/(1-f))$, where $W(z)$ is the Lambert W function. Expanding for small $f$:

$$f \approx \sqrt{1+2(1+\phi)\lambda/d} - 1 \approx (1+\phi)\lambda/d. \tag{4}$$

For $\lambda/d \ll 1$, compaction is linear in $\lambda/d$ and loop nesting is unimportant because it is rare (Fig. 3(b) in the main text).

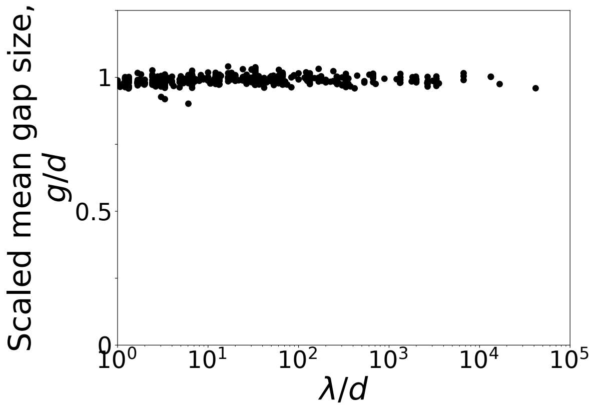


FIG. 1. In simulations, we observe scaled mean gap size $g/d \approx 1$ in the dense ($\lambda/d > 1$) regime, in agreement with the mean-field theory.

*
†
